# Reconfiguring primase DNA-recognition sequences by using a data-driven approach

**DOI:** 10.1101/2020.09.29.317842

**Authors:** Adam Soffer, Morya Ifrach, Stefan Ilic, Ariel Afek, Hallel Schussheim, Dan Vilenchik, Barak Akabayov

## Abstract

DNA-protein interactions are essential in all aspects of every living cell. Understanding of how features embedded in the DNA sequence affect specific interactions with proteins is challenging but important, since it may contribute to finding the means to regulate metabolic pathways involving DNA-protein interactions. Using a massive experimental benchmark dataset of binding scores for DNA sequences and a machine learning workflow, we describe the binding to DNA of T7 primase, as a model system for specific DNA-protein interactions. Effective binding of T7 primase to its specific DNA recognition sequences triggers the formation of RNA primers that serve as Okazaki fragment start sites during DNA replication.

## INTRODUCTION

Specific protein-DNA recognition is essential for a wide range of cellular processes, including DNA replication, repair, and recombination (1). Determination of the specific binding preferences of proteins both *in vivo* (2–7) and *in* vitro (8–17) has been facilitated by recent technological advances in high-throughput testing. Computational analysis combined with such high-throughput assays have identified protein-DNA binding preferences on the whole-genome level (8,11,16,18-25), and the information so obtained has been used to elucidate mechanisms of the gene expression regulation by transcription factors (TFs) and RNA polymerases in different organisms (20,24,26,27).

DNA replication serves as a metabolic pathway in all cells in which specific DNA-protein interactions take place(28). During DNA replication the double-stranded DNA is unwound to expose the two individual DNA strands; one is copied continuously (leading strand) and the other is copied discontinuously (lagging strand). On the lagging DNA strand, an enzyme known as DNA primase recognizes the DNA sequence used as the template for the synthesis of RNA primers, and DNA polymerase is then responsible for elongating these RNA primers into the DNA segments known as Okazaki fragments. This process of RNA-primed DNA synthesis by a DNA polymerase is triggered exclusively by the recognition of a specific DNA sequence by the primase. This recognition is thus fundamental to the establishment of Okazaki fragments and consequently to the whole process of proper DNA replication. In prokaryotes, RNA primer formation occurs on pre-defined sequences on the genome that are specifically recognized by a DnaG-type primase (29) (Figure 1). For example, the activity of bacteriophage T7 DNA primase, comprising the N-domain [residues 1-271] of gene 4 protein, is initiated by sequence-specific binding of DNA primase to 5’-GTC-3’ (30,31), which is then followed by the synthesis of a functional primer (32). Importantly, it is now known that even though DNA primase recognizes a specific trinucleotide sequence, flexibility in the selection of initiation sites for Okazaki fragments is allowed (33), i.e., not every primase-DNA recognition sequence (PDRS) will become an Okazaki fragment start site. The reason for this flexibility is, however, not understood and is the enigma that we address in the current study: Although extensive research has been carried out on the interactions of DNA primase with DNA, it is still not clear why DNA primase ignores the majority of trinucleotide recognition sites. In *Escherichia coli*, for example, DnaG primase ignores ~97% of the trinucleotide recognition sites and initiates Okazaki fragments only every 1.5-2 kb (and not more frequently)(34). The literature offers two possible explanations for the effect of selective DNA sequence recognition by a primase: The first, well-explored possibility, is that other DNA replication proteins, such as DnaB (35–38), single-stranded DNA-binding protein (SSB)(39), or clamp loader (40), affect the binding of the primase (DnaG) to DNA or even change the preferences for PDRSs on the genome. The second possibility is that a sequence larger than a trinucleotide is required for the specific binding of a primase. On the basis of the mismatch between the frequency of GTC sequences on the bacteriophage T7 genome and the actual size of the Okazaki fragments, it is reasonable to assume that only a sequence larger than a trinucleotide will lead to “effective binding” of a DNA primase, i.e., binding yielding an RNA primer that marks the start site of an Okazaki fragment.

**Figure 1.**
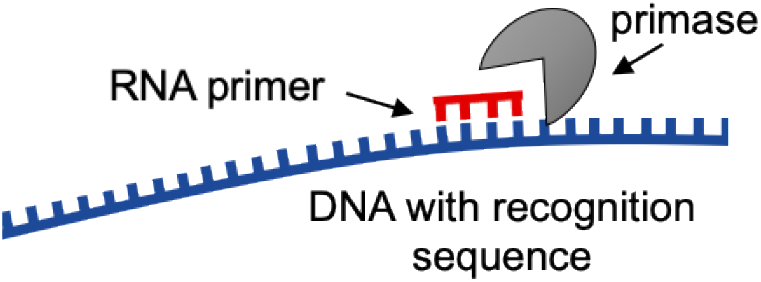
Schematic representation of primase binding to a single-stranded DNA template and synthesis of an RNA primer.

In our first steps to investigate the above-described possibility of a larger DNA binding determinant, we applied high-throughput primase profiling (41), in which binding scores (referred as median fluorescence intensities) for tens of thousands of DNA primase–DNA binding events on a protein-DNA binding microarray (PBM) were combined with biochemical analysis (41). This technology facilitated the analysis of the composition of sequences flanking the specific recognition site and their corresponding binding scores and confirmed that a GTC sequence is not sufficient for the recognition of DNA templates by T7 primase (41,42). Specifically, we showed that T7 primase has a high affinity for PDRSs composed of GTC with G/C/T-rich flanking sequences, leading to the formation of longer RNA primers (41). Although the development of high-throughput primase profiling facilitated the acquisition of massive amounts of data on DNA-primase-binding events, the means to systematically analyze that data in a way that will facilitate a comprehensive understanding of the recognition process are yet to be put in place. Specifically, analysis of this data will throw light on the principles for selection of PDRSs on a genome during DNA replication by enabling us to answer the following questions: 1) Is there information stored in the DNA sequence that is important for T7 primase binding? 2) If we understand the principles of specific primase DNA recognition, can we predict binding scores of T7 primase for a given DNA sequence? And 3) Can we generate new DNA sequences with desired binding scores based on the sequence features embedded in the DNA? Answering the third question will enable us to ascertain which features embedded in specific DNA binding sequences govern the binding of DNA primase.

In concert with cutting-edge developments in biochemical technologies, current progress in computational science provides us with the opportunity to construct knowledge-based models that will help us to answer the above questions. Here, we describe an intelligent learning workflow that provides a comprehensive view of the principles that govern the design and activity of PDRSs with unprecedented flexibility and accuracy. We applied this workflow to elucidate the link between the larger context, i.e., the flanking nucleotides, of the primase recognition sequence and the synthesis of RNA primers that initiate Okazaki fragments. Whereas data in Afek at al shows that TA is better than GA, here we found determinants and set of rules that allow us to predict binding and catalysis of DNA primase. Specifically, using state-of-the-art machine learning analysis, we found set of rules, that allowed quantitative prediction of primase binding for any DNA sequence.

## MATERIAL AND METHODS

Detailed descriptions of the materials and methods used in this work, including design of DNA library, data-preprocessing, machine learning algorithms (unsupervised and supervised), protein expression/purification, and DNA primase activity, are provided in SI Appendix, Materials and Methods.

## RESULTS AND DISCUSSION

The overall structure of our study is comprised of the following stages of analysis of the data obtained using PBMs for quantitative measurements of T7 primase-DNA binding, after preprocessing of the data (Scheme 1): <u>step 1</u>) preparation of a PBM-driven benchmark data; <u>step 2</u>) clustering the PDRSs containing DNA sequences; <u>step 3</u>) training a regression model; and <u>step 4</u>) predicting the score of new DNA sequences, and generating novel DNA sequences with desired binding scores for T7 primase. These steps are elaborated below, as are the data preprocessing and <u>step 5</u> (Scheme 1), which is biochemical validation.

### Data preprocessing and vectorization of DNA sequences

Data acquisition was obtained using His-tagged T7 primase produced and purified as described previously (41). All algorithms described for data preprocessing and analysis are written in python and are publicly available (https://github.com/csbarak/T7pdrs). Before the data analysis, considerable attention was paid to data preprocessing, as the success of the subsequent application of machine-learning algorithms depended on the explicit presentation of the data in a way that facilitated the extraction of meaningful features and the removal of distracting outliers. The preprocessing of PBM-derived DNA-primase binding data comprised four steps: data cleansing, data filtration, embedding of the sequences into vectors, and data normalization, as follows. Using the PDRSs as meaningful “words” on the basis of their sequence features, where each sequence was assigned to its PBM-driven binding score, we focused on the sequences that could potentially serve as Okazaki fragment start sites. We started with the preparation of a “lexicon” of DNA “words,” each comprising a larger context of GTC-containing sequences that allow effective binding of T7 primase, i.e., the binding of T7 primase that yields RNA primers. The starting point for the preprocessing was that while an average size of ~64 nucleotides (Figure 2a) is the expected distance between two GTC sequences, Okazaki fragment size of 1000-6000 nucleotides is obtained experimentally (Figure 2b, marked in the red range box).

**Figure 2.**
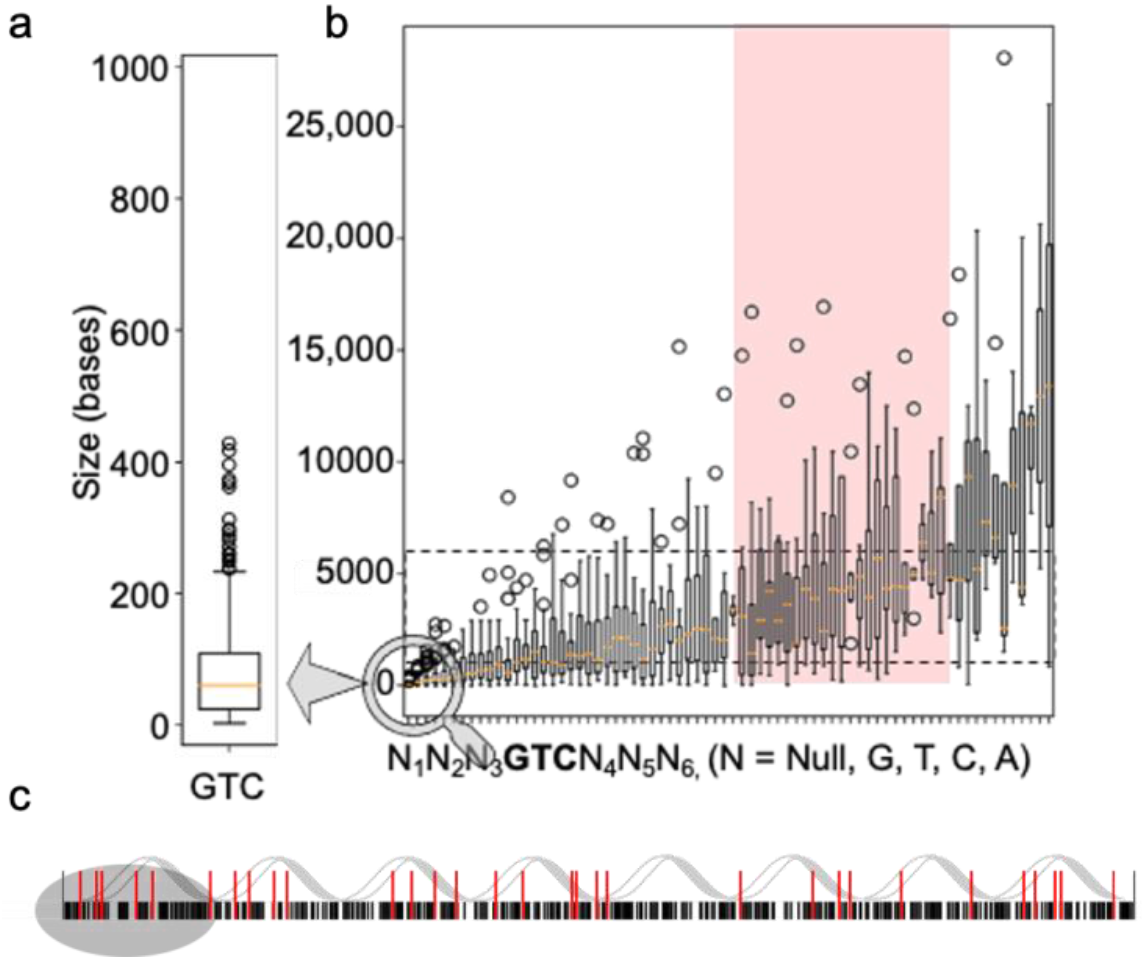
Frequency and distribution of GTC-containing primase-DNA recognition sequences (PDRSs) on the bacteriophage T7 genome. a, The frequency of the occurrence of GTC in a random sequence is every 4^3^ = 64 bases (approximately as indicated by the orange line). b, Calculated size distribution of the DNA sequence between GTC-containing PDRSs on the T7 genome that match the actual size of Okazaki fragments. These PDRSs consist of 0-3 nucleotides flanking the GTC sequence and are distributed at an inter-PDRS distance that ranges between 1000-6000 nucleotides, which yield Okazaki fragments of the same size. c, All combinations of possible T7 PDRSs (5’-GTC-3’) on the genome are considered. Black lines represent the frequency of GTCs; red lines represent the frequency of large-context GTC-containing PDRSs that match experimental values for Okazaki fragment sizes.

We thus posited that GTC-containing PDRSs must be larger than a trinucleotide sequence to meet the frequency on the genome that would allow the creation of Okazaki fragments of sizes that were observed previously (Figure 2c, red lines: larger GTC-containing sequences that match experimental sizes of Okazaki fragments (43); black lines: frequency of GTC every 64 bases).

Since DNA sequences constitute a form of categorical data represented by nucleotides, the preprocessing step was required to convert the plain representation of DNA sequences into a meaningful numeric representation. Such a representation of DNA sequences was obtained by using One Hot Encoding (OHE). In this way, a categorical sequence was converted into an array of integers in which each nucleotide was represented by four unit vectors: (A=[1000], C=[0100], G=[0010], T=[0001]), e.g., the sequence ACCG was encoded as 1000|01000|01000|0010. Every DNA sequence, represented by a 144-dimensional vector, was fed as an input into both a Kmeans model (using Euclidian distance) and a Ward-method-based (44) hierarchical clustering model.

### Defining the mathematical descriptors (features) of the PDRSs

The challenge in the selection of descriptors in the DNA sequences derived from the fact that only a limited number of features that have chemical/physical meaning are useful for model construction and from difficulties in converting DNA sequences into vectors of numbers. Importantly, nucleotides, being categorical variables, cannot be treated in terms of ordinal data. Since “hand-crafted” features extracted from DNA sequences did not improve prediction of primase binding scores (Supplementary Figure S1), we utilized the K-mer method for feature extraction (45). In brief, the K-mer is a frequency vector that counts all possible combinations of short sequences (of size K) in larger DNA sequences. As the K parameter increases, the number of possible combinations increases, while the frequency of each mer decreases (Supplementary Figure S2), giving sparser, yet more detailed, data. Since the K-mer method can be implemented with different normalization and striding techniques, even with insufficient structural information, we used a 1-step stride to allow overlap between mers and normalized the extracted K-mer counts.

Extracting features from the DNA sequences of the microarray using the K-mer approach allowed us to find association rules for those DNA sequences (unsupervised algorithms). The sequential features obtained were also used to generate a prediction model of primase-DNA binding, based on primase-binding data collected from PBM experiments (supervised algorithms).

### Exploratory data analysis

As is customary, we started with exploratory data analysis, which is unsupervised in nature (i.e., the primase binding scores were ignored). The goal here was to produce a meaningful visualization of the data with the aim to obtain new insights. To this end, we reduced the dimensionality of the data using principal component analysis (PCA), and applied various clustering algorithms, which revealed a meaningful cluster structure with respect to the binding score. After applying PCA to the data, i.e., to the 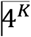 dimensions existing for each 36-mer oligonucleotide, we found that the top three principal components explained 64% of the total variance. Therefore, we concluded that this number of principal components enables the production of a meaningful 3D picture.

To interpret the clusters generated using PCA, the binding values obtained for the T7 primase of all the DNA sequences were normalized and used to color code the data points in the clusters (Figure 3a). The most striking result to emerge from the color-coded data was its arrangement into five clusters—one homogeneous cluster of DNA sequences that are strongly bound to T7 primase (colored red in Figure 3a), two homogeneous clusters of DNA sequences with weaker binding to T7 primase (colored blue), and two inhomogeneous clusters with uniformly distributed binding scores. This organization of the data points into meaningful clusters indicates that: 1) there are hidden descriptors within the DNA sequence that are essential for primase binding, and 2) the sequence descriptors obtained by the K-mer approach (Figure 3b) are more than adequate for describing primase–DNA binding.

**Figure 3.**
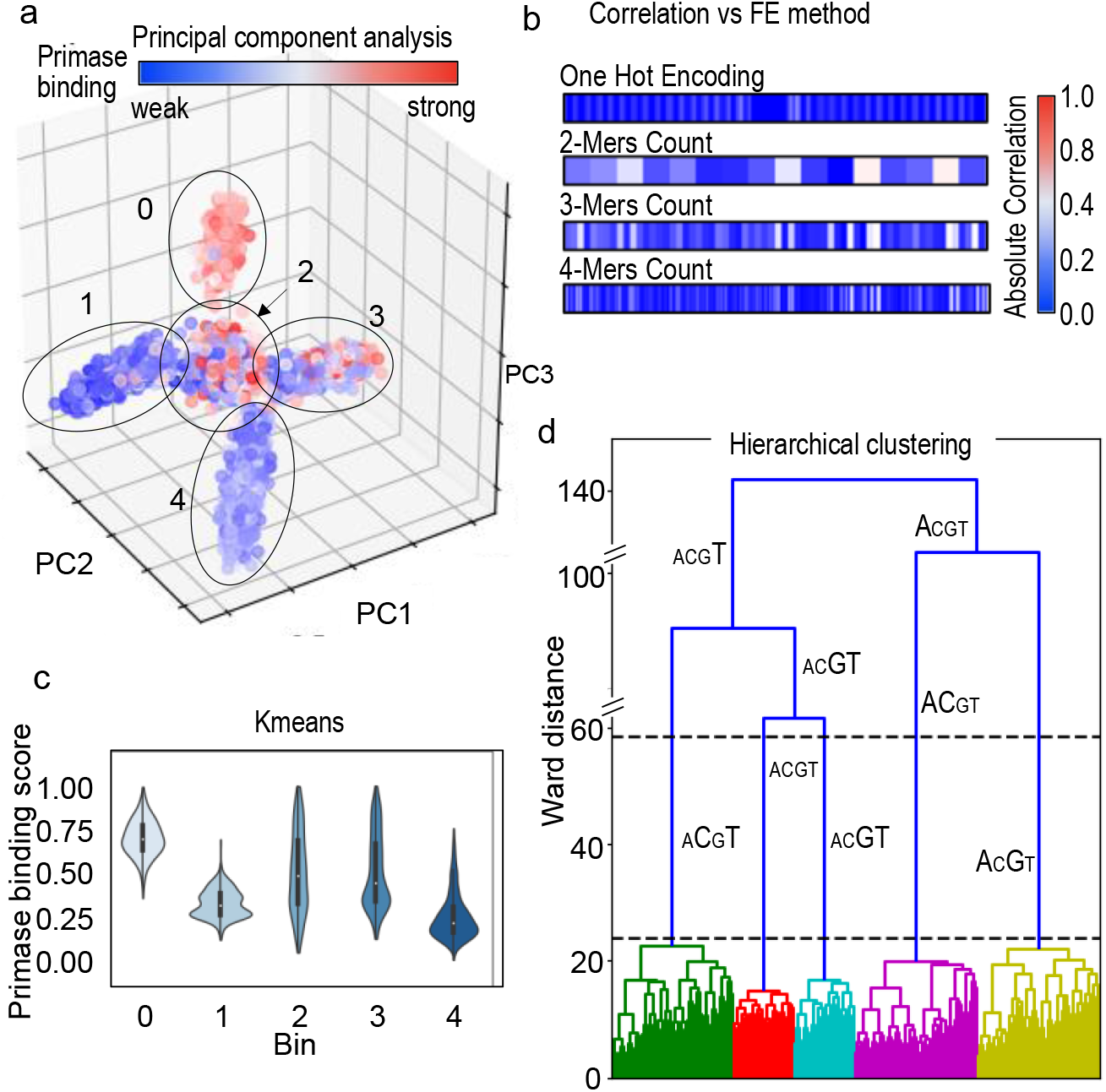
Inference from DNA sequences without labeled responses for T7 primase binding (unsupervised learning). a, Dimensionality reduction algorithm PCA tri-plot used to visualize the data by selecting the top three principal components. Assignment of the binding scores (labels) to each data point shows an uneven distribution across two clusters (2, 3) and a homogeneous distribution in three clusters (0, for strong binding to T7 primase, and 1 and 4 for weak binding). b, Correlation between the primase binding score and feature extraction (FE) using different methods: OHE, 2-mer, 3-mer and 4-mer counts. K-mers were used as descriptors for the PCA analysis. c, Kmeans clustering on one-hot encoded DNA sequences. Clustering was performed by measuring pairwise distances of DNA sequences from the centroid of each cluster. Violin plots representing the distribution of the binding scores assigned to each data point in the clusters are shown. Three clusters show evenly distributed scores (0, 1,4) and two show a less homogeneous score distribution (2, 3). Each cluster is represented by a “mean word” (centroid): (0) GTTTTGTTTTGTTTTTGTC GTGTGGTTGTGGTGGTA; (1) CTTTTTTTCTTTTTTCGTCCTTTTTT TTTTTCCCCA; (2) GAAGAAAATCCATAGGGTCAACCGGGTTATG TTAAA; (3) CCACAAAAAAAAAAAAGTCCAACCCACAAAACCCC A; (4) GGAAAGGAGAGAAAAAGTCAAAAAAGAAAGAAAGAA d, Hierarchical clustering. The x-axis shows DNA sequences emerging into clusters, and the y-axis shows the induced Ward distance of each stage. Letter sizes indicate the letter’s frequency in each cluster. The maximal Ward distance gap is indicated between the dashed black lines. The figures were created using the Python package Seaborn and Matplotlib.

Using the Kmeans algorithm, we were able to shed light on the distributions of the binding scores within the five clusters and to show that preprocessing using OHE results in clusters containing similar score distributions to the 5-cluster structure obtained by PCA. The Kmeans iterative algorithm partitions the data space into sub-spaces, thereby assigning a matching label (the cluster number) to each instance according to its location.

Clustering of the unlabeled DNA sequences in Kmeans relied on the sequence distances from the corresponding cluster centroids. As each sequence was represented using OHE, the distance between two sequences could be described as the number of changes needed in one sequence to convert it into the other sequence. In the Kmeans analysis, aligning the PBM-driven binding score for each DNA sequence revealed that each cluster exhibited a clear trend, as the group of binding scores to the primase was distributed unevenly with long tails (Figure 3c), which means that each cluster held exceptions.

While Kmeans computes distances of instances from the centroids, clustering of unlabeled DNA sequences using Ward’s minimum variance method (44) allowed tracking of the evolution of clusters (Figure 3d). The maximal Ward distance (WD) gap was obtained using 5 clusters, as observed from PCA and Kmeans for the same dataset of DNA sequences. Furthermore, we can see in Figure 3d that each colored branch holds a sub-group with two highly repeating letters (CT, GT, AC, AG) or a uniform distribution of the letters (ACGT).

### Predicting the binding score

After clustering the DNA sequences in the microarray into groups with common features, the next step was to predict the outcome of T7 primase–DNA binding for a given DNA sequence. To increase accuracy, each cluster was fitted separately. We used GTC-containing DNA sequences and their corresponding PBM-driven binding scores as input and output pairs for the training set, respectively. The PBM-driven data comprises the continuous numerical binding values for DNA sequences, i.e., the type of data that regression models are aimed to solve.

We extracted sequence-based features (SBF) inspired by pseudo K-tuple nucleotide composition (46). We modified the ordinary method for SBF extraction by neglecting locality-based features, since distance was interpreted as the number of nucleotides between a mer and its closest 5’-GTC start site. Our modified method for feature extraction provided us with sequence-wise normalized K-mers (*see* Methods for normalization of the K-mers). We then tested different regression algorithms using criteria that can differentiate between uninformative and highly informative SBFs. Thereafter, we applied the L1 regularized linear regression [least absolute shrinkage and selection operator (Lasso) (47)] model on each bin separately with the aim to emphasize meaningful mers and to prevent overfitting of the model. (Lasso’s output is a closed form expression that is generated under fewer, yet more meaningful, coefficients by applying a penalty for each variable.) In addition, we extracted an expected performance measure separately for each bin obtained by Kmeans, as the mean absolute errors (MAE, an error estimate for the regression) for our results were uneven across bins (Table 1). As we trained five Lasso models on 5 bins, each bin’s prediction was obtained using a different set of coefficients. While overall performance of the prediction model is obvious, examining the performance for each bin is more precise. Lasso’s performance differed across bins, where bins 0 and 1 each generated an error of about 15%, and bins 2, 3 and 4 generated a relatively small error of 6% (Table 1). We found that pre-treating the data using Kmeans decreased the prediction error by approximately 10% (Table 2).

**Table 1.** Results summary of 5-Fold-MCCV* in each clusters

| Bin | MAE* | STD | Mean MAE [%] | Mean STD [%] | Bin Weight [% of instances from data] |
| --- | --- | --- | --- | --- | --- |
| 0 | 0.168 | 0.005 | 16.8 | 0.5 | 15% |
| 1 | 0.138 | 0.007 | 13.8 | 0.7 | 19.4% |
| 2 | 0.067 | 0.004 | 6.7 | 0.4 | 13.27% |
| 3 | 0.060 | 0.003 | 6 | 0.3 | 25.3% |
| 4 | 0.077 | 0.003 | 7 | 0.3 | 26.8% |

\*MAE, mean absolute error; MCCV, Monte Carlo cross-validation

**Table 2.** Effect of Kmers on model prediction

| MAE* | K=1 | K=2 | K=3 | K=4 |
| --- | --- | --- | --- | --- |
| BNS = 1 | 0.171 | 0.112 | 0.103 | 0.084 |
| BNS = 5 | 0.110 | 0.102 | 0.093 | 0.079 |
| Ratio $\frac{\text{error no bins}}{\text{error with bins}}$ | 1.55 | 1.09 | 1.10 | 1.06 |
| Decrease in error in %<br>$100 * \left(1 - \frac{\text{error with bins}}{\text{error no bins}}\right)$ | 35% | 9% | 10% | 6% |

\*MAE, mean absolute error

To force all the coefficients of the Lasso model to be positive for every K-mer feature, we extracted the largest 10 coefficients of each bin and investigated the effect of the K-mer features on the model. The trained models were cross validated on 5-folds of the training dataset and tested on a small test-set taken from the PBM results; the set consisted of 16 sequences, divided into two groups with significantly different primase binding signals (Figure 4a), namely, 8 sequences that showed weak binding to primase, and 8 sequences with strong binding. Given the training data distribution, our models predicted binding of primase with any GTC-containing sequence, with an MAE of < 12% (Figure 4).

**Figure 4.**
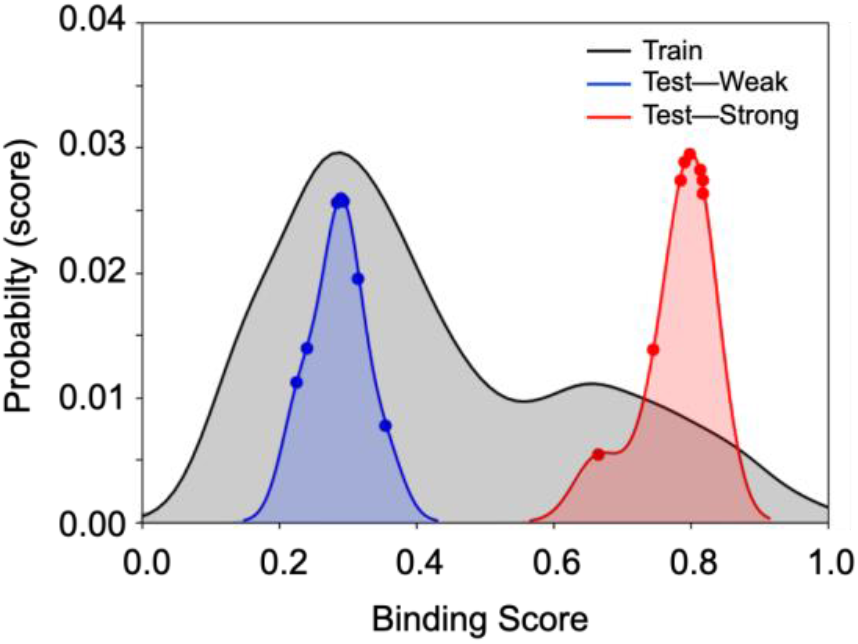
Results for linear regression with L1 regularization (Lasso). After cleaning, the training set contained 3150 instances (DNA sequences), whereas the test set contained 16 instances. Prediction of scores by using the regression model was performed on 16 DNA sequences with known scores, eight of which showed weak binding to T7 primase (blue graph) and eight showed strong binding to T7 primase (red graph). In accordance with the training-set double distribution (black graph), the predicted binding of the two test groups are distributed at weak and strong binding scores areas, respectively. Although the probability of finding DNA sequences with strong binding to primase is low, the model accurately predicted all DNA sequences that belong to the strong binding group. DNA sequences, their empirical and predicted scores are presented in Supplementary Table S1.

In order to identify the most influential nucleotides for accurate prediction of binding score for a given PDRS, we used a different perspective on the data. Each nucleotide position before and after the GTC sequence was referred as a feature and trained against Gradient Boosting Machine (GBM) and Random Forest models. GBM and Random Forest models can handle categorical data, as DNA sequences, and can easily explore the importance of each feature for the model. The results indicate that adjacent nucleotides to the GTC sequence are the most essential for primase binding, whereas distant nucleotides are much less influential (Supplementary Figure S3).

### Biochemical validation

On the genome, the initial step of sequence specific (PDRS) binding is followed by synthesis of a dinucleotide (5’-AC-3’), which is then extended into a functional primer by DNA primase. It has previously been shown that A/G-containing sequences that flank the specific recognition site increase primase-DNA binding affinity in comparison to T/G-containing sequences^14^. Since binding to DNA is a pre-requisite for primase activity, the strength of binding affects the magnitude of the catalytic activity of the primase and the yield of the RNA product. We used qualitative biochemical assays to experimentally validate the prediction model described above. The validation provided insights into the features embedded in the DNA sequences that are important for binding and catalytic activity of DNA primase.

#### I. Correlation between prediction of primase binding to PDRS and catalytic activity

The eight sequences with strong binding and the eight with weak binding to primase, which were used as the test set in the supervised learning part of this study, yielded RNA primers, as was expected from their PBM-driven binding values. This finding shows, for the first time, that the sequence descriptors embedded in the DNA sequence are sufficient to predict binding scores and that prediction of a binding sequence correlates well (96.9 % Pearson correlation coefficient) with the formation of RNA primers (Supplementary Figure S4). The understanding of how sequential features embedded in the DNA is related to the binding of primase allows us not only to predict binding scores of a given PDRS, but also to design novel PDRSs that yield high primase binding scores.

#### II. Exhaustive search for flanking sequences that yield novel PDRSs

Features important for DNA-primase binding were used in formulating design principles to generate novel GTC-containing DNA sequences with desired binding scores. Assuming that the DNA sequences originate from 5 different clusters (Figure 3) that require 5 different models, we generated two types of DNA sequences as follows: 1) We selected DNA sequences from two homogenous clusters of primase binding scores and exhaustively altered the non-”GTC” nucleotides to generate primase recognition DNA sequences (new PDRSs). Three altered sequences that did not exist in the training set and yielded the 10th, 50th, and 90th percentile binding strengths were selected from the two clusters (clusters 0 and 4, Figure 3) for further biochemical evaluation. 2) DNA sequences were generated in the same way, and two novel DNA sequences from each Kmean cluster, one that represented the strongest binding prediction and another that represented weakest, were selected (overall 10 sequences. DNA sequence and their predicted scores are presented in Supplementary Table S2) for biochemical analysis (Supplementary Figure S5).

To characterize the effect of the novel DNA sequences (PDRSs) generated as described above, we quantified and compared RNA primer formation by T7 primase, where the generated PDRSs were used as templates for the synthesis (Figure 5). Specifically, we used [γ–^32^P]ATP to 5’ end-label the RNA primers, which ensured that each primer was labeled only once, and thus the absolute amounts of RNA primers could be quantified (Figure 5). For the 10 DNA templates that represented weak/strong binding to primase from each one of the five clusters (Figure 3), we found that the newly designed DNA sequence flanking the 5’-GTC-3’ sequence with higher binding scores for T7 primase showed improved RNA primer synthesis activity, as was to be expected (Supplementary Figure S5). These results confirm our machine learning prediction model and indicate that higher binding affinity for PDRS recognition sequences are dictated by features embedded in the DNA sequence. Using the <u>exhaustive search for flanking sequences that yield novel primase-DNA binding sequences developed here</u>, we are now in a position to design DNA templates that yield: 1) larger amounts of RNA primers, and 2) longer RNA primers that can serve as functional primers for T7 DNA polymerase(48). Both the length and the quantity of the RNA primers are likely to be essential for the decision to start Okazaki fragments by DNA polymerase on the lagging DNA strand.

**Figure 5.**
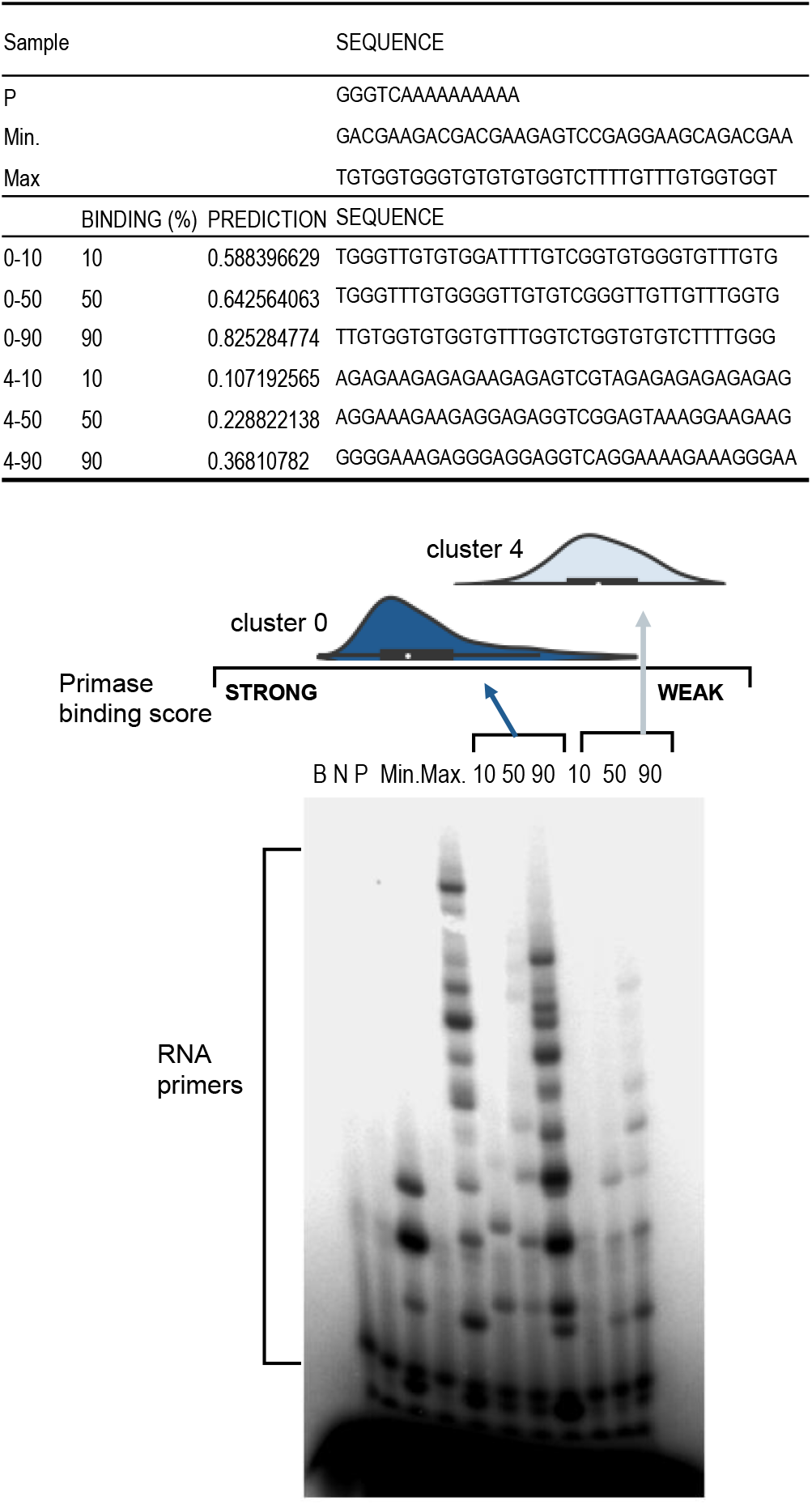
RNA primer synthesis catalyzed by the T7 primase on computer generated GTC-containing DNA templates. Three DNA sequences from Kmeans clusters #0 and #4 that predicted the 10th, 50th, and 90th percentile binding scores were selected in each cluster. Distribution of binding values for the two clusters are presented. Note that cluster #0 shows stronger primase binding values, on average, than cluster #4. The standard reaction mixture contained oligonucleotides with the primase recognition sequence, a control oligonucleotide 5’-GGGTCA10-3’, and γ-^32^P-ATP, CTP, GTP, and UTP. After incubation, the radioactive products were analyzed by electrophoresis on a 25% polyacrylamide gel containing 7 M urea, and visualized using autoradiography. Table present the DNA sequences and their corresponding binding values.

## DISCUSSION

On the basis of PBM data for primase binding previously obtained for > 150,000 DNA sequences, this study set out to: 1) develop the means to predict the binding score of T7 primase for a given DNA sequence, 2) describe the DNA sequence features essential for binding of the enzyme, and 3) generate novel sequences with a high propensity for T7 primase binding. The K-mers approach for feature selection in the DNA sequential data that was used here appears to cover all possible combinatorial pieces of information hidden in the DNA sequences and serves as an efficient strategy for feature extraction. The Kmers method, which simply counts explicit combinations of nucleotides in a DNA sequence, was superior to other accepted methods of “hand-crafted” feature extraction from DNA sequences. Features obtained by Kmers clearly bear DNA properties important for primase binding, as demonstrated by the unsupervised analysis in which clustered groups coincided with experimental binding scores.

Although this study focused on DNA sequence recognition by T7 primase, the findings may well have bearing on rules hidden in DNA sequences that are crucial for other specific DNA-protein interactions. These findings thus contribute to our understanding of how DNA primase selects Okazaki fragments start sites on the genome and why only some of the possible priming sites initiate during DNA replication, while others do not, resulting in Okazaki fragments with a larger-than-expected average length. The implications of this study are that design principles for any DNA sequence with a desired binding affinity to T7 primase can indeed be generated computationally on the basis of our analysis. Furthermore, PDRSs could be designed to yield an RNA primer with a particular content. In conclusion, state-of-the-art carefully selected learning methods, like those used here, have enormous analytical potential for predicting specific protein-DNA interactions, but require large amounts of data, a requirement than can indeed be met by using PBMs. Our results for T7 DNA primase as a model system, can generalize to other primases with improved sensitivity and specificity.

## DATA AVAILABILITY

All datasets and python codes for algorithms used for preprocessing and analysis are available in the GitHub repository (https://github.com/csbarak/T7pdrs).

## SUPPLEMENTARY DATA

Experimental and computational methods, supplementary tables and figures. Supplementary Data are available at NAR online.

## FUNDING

This research was supported by the Israel Science Foundation (ISF, Grant No. 1023/18).

## CONFLICT OF INTEREST

The authors declare that there is no conflict of interest.

---

**Scheme 1.**
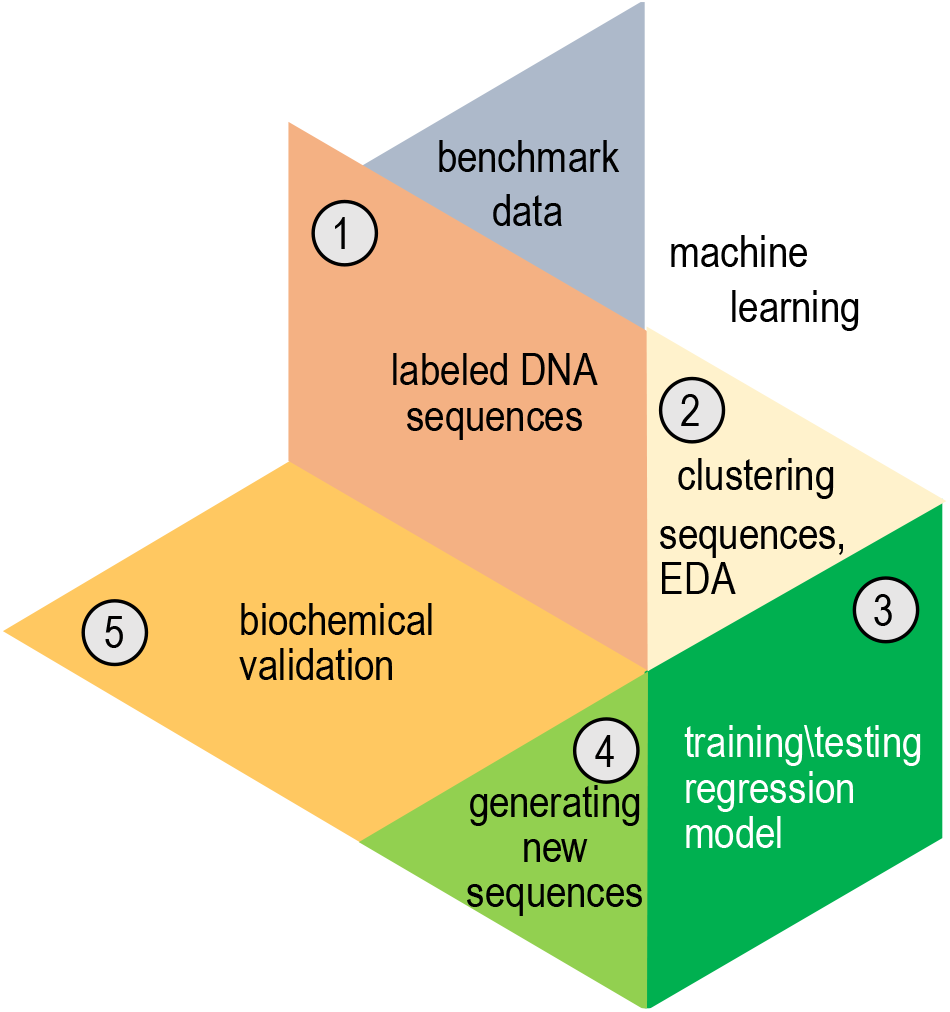
Analysis workflow after preprocessing the data from the primase-DNA binding microarray. The benchmark dataset containing DNA sequences for the training set was preprocessed (step 1). The DNA sequences were clustered into five bins using exploratory data analysis (EDA), i.e., unsupervised algorithms (step 2). A different regressor was trained for every cluster. Several regression algorithms were used; linear regression with L1 regularization provided the best results. To predict the binding scores of a new DNA sequence, the sequence was assigned to a specific bin, and its score for primase binding was predicted using that bin’s regressor (step 3). Novel DNA sequences (PDRSs) with high binding score for primase were generated (step 4). It was then possible to examine the ability of those PDRSs to bind primase and induce the synthesis of RNA primers (step 5).

## Notes

### Competing Interest Statement

The authors have declared no competing interest.

### Summary of Updates

The title abstract and the first paragraph of the introduction have been slightly modified. Non of the results/discussion/conclusions were modified.

## REFERENCES

1. Lodish, H.F., Berk, A., Zipursky, S.L., Matsudaira, P., Baltimore, D. and Darnell, J. (2000) Molecular cell biology. Citeseer.

2. Boyle, A.P., Davis, S., Shulha, H.P., Meltzer, P., Margulies, E.H., Weng, Z., Furey, T.S. and Crawford, G.E. (2008) High-resolution mapping and characterization of open chromatin across the genome. Cell, 132, 311–322.

3. Giresi, P.G., Kim, J., McDaniell, R.M., Iyer, V.R. and Lieb, J.D. (2007) FAIRE (Formaldehyde-Assisted Isolation of Regulatory Elements) isolates active regulatory elements from human chromatin. Genome research, 17, 877–885.

4. Hesselberth, J.R., Chen, X., Zhang, Z., Sabo, P.J., Sandstrom, R., Reynolds, A.P., Thurman, R. E., Neph, S., Kuehn, M.S. and Noble, W.S. (2009) Global mapping of protein-DNA interactions in vivo by digital genomic footprinting. Nature methods, 6, 283–289.

5. Johnson, D.S., Mortazavi, A., Myers, R.M. and Wold, B. (2007) Genome-wide mapping of in vivo protein-DNA interactions. Science, 316, 1497–1502.

6. Ren, B., Robert, F., Wyrick, J.J., Aparicio, O., Jennings, E.G., Simon, I., Zeitlinger, J., Schreiber, J., Hannett, N. and Kanin, E. (2000) Genome-wide location and function of DNA binding proteins. Science, 290, 2306–2309.

7. Rhee, H.S. and Pugh, B.F. (2011) Comprehensive genome-wide protein-DNA interactions detected at single-nucleotide resolution. Cell, 147, 1408–1419.

8. Badis, G., Berger, M.F., Philippakis, A.A., Talukder, S., Gehrke, A.R., Jaeger, S.A., Chan, E.T., Metzler, G., Vedenko, A. and Chen, X. (2009) Diversity and complexity in DNA recognition by transcription factors. Science, 324, 1720–1723.

9. Berger, M.F., Philippakis, A.A., Qureshi, A.M., He, F.S., Estep, P.W. and Bulyk, M.L. (2006) Compact, universal DNA microarrays to comprehensively determine transcription-factor binding site specificities. Nature biotechnology, 24, 1429–1435.

10. Fordyce, P.M., Gerber, D., Tran, D., Zheng, J., Li, H., DeRisi, J.L. and Quake, S.R. (2010) De novo identification and biophysical characterization of transcription-factor binding sites with microfluidic affinity analysis. Nature biotechnology, 28, 970–975.

11. Jolma, A., Yan, J., Whitington, T., Toivonen, J., Nitta, K.R., Rastas, P., Morgunova, E., Enge, M., Taipale, M. and Wei, G. (2013) DNA-binding specificities of human transcription factors. Cell, 152, 327–339.

12. Maerkl, S.J. and Quake, S.R. (2007) A systems approach to measuring the binding energy landscapes of transcription factors. Science, 315, 233–237.

13. Noyes, M.B., Christensen, R.G., Wakabayashi, A., Stormo, G.D., Brodsky, M.H. and Wolfe, S. A. (2008) Analysis of homeodomain specificities allows the family-wide prediction of preferred recognition sites. Cell, 133, 1277–1289.

14. Riley, T.R., Slattery, M., Abe, N., Rastogi, C., Liu, D., Mann, R.S. and Bussemaker, H.J. (2014) SELEX-seq: a method for characterizing the complete repertoire of binding site preferences for transcription factor complexes. Hox Genes: Methods and Protocols, 255–278.

15. Warren, C.L., Kratochvil, N.C., Hauschild, K.E., Foister, S., Brezinski, M.L., Dervan, P.B., Phillips, G.N. and Ansari, A.Z. (2006) Defining the sequence-recognition profile of DNA-binding molecules. Proceedings of the National Academy of Sciences of the United States of America, 103, 867–872.

16. Weirauch, M.T., Yang, A., Albu, M., Cote, A.G., Montenegro-Montero, A., Drewe, P., Najafabadi, H.S., Lambert, S.A., Mann, I. and Cook, K. (2014) Determination and inference of eukaryotic transcription factor sequence specificity. Cell, 158, 1431–1443.

17. Zykovich, A., Korf, I. and Segal, D.J. (2009) Bind-n-Seq: high-throughput analysis of in vitro protein–DNA interactions using massively parallel sequencing. Nucleic acids research, gkp802.

18. Birney, E., Stamatoyannopoulos, J.A., Dutta, A., Guigó, R., Gingeras, T.R., Margulies, E.H., Weng, Z., Snyder, M., Dermitzakis, E.T. and Thurman, R.E. (2007) Identification and analysis of functional elements in 1% of the human genome by the ENCODE pilot project. Nature, 447, 799–816.

19. Carlson, C.D., Warren, C.L., Hauschild, K.E., Ozers, M.S., Qadir, N., Bhimsaria, D., Lee, Y., Cerrina, F. and Ansari, A.Z. (2010) Specificity landscapes of DNA binding molecules elucidate biological function. Proceedings of the National Academy of Sciences, 107, 4544–4549.

20. Consortium, E.P. (2004) The ENCODE (ENCyclopedia of DNA elements) project. Science, 306, 636–640.

21. Hume, M.A., Barrera, L.A., Gisselbrecht, S.S. and Bulyk, M.L. (2014) UniPROBE, update 2015: new tools and content for the online database of protein-binding microarray data on protein–DNA interactions. Nucleic acids research, gku1045.

22. Lee, T.I., Rinaldi, N.J., Robert, F., Odom, D.T., Bar-Joseph, Z., Gerber, G.K., Hannett, N.M., Harbison, C.T., Thompson, C.M. and Simon, I. (2002) Transcriptional regulatory networks in Saccharomyces cerevisiae. Science, 298, 799–804.

23. Rohs, R., West, S.M., Sosinsky, A., Liu, P., Mann, R.S. and Honig, B. (2009) The role of DNA shape in protein-DNA recognition. Nature, 461, 1248–1253.

24. Roy, S., Ernst, J., Kharchenko, P.V., Kheradpour, P., Negre, N., Eaton, M.L., Landolin, J.M., Bristow, C.A., Ma, L. and Lin, M.F. (2010) Identification of functional elements and regulatory circuits by Drosophila modENCODE. Science, 330, 1787–1797.

25. Zhao, Y. and Stormo, G.D. (2011) Quantitative analysis demonstrates most transcription factors require only simple models of specificity. Nature biotechnology, 29, 480–483.

26. Mooney, R.A., Davis, S.E., Peters, J.M., Rowland, J.L., Ansari, A.Z. and Landick, R. (2009) Regulator trafficking on bacterial transcription units in vivo. Molecular cell, 33, 97–108.

27. Venters, B.J. and Pugh, B.F. (2013) Genomic organization of human transcription initiation complexes. Nature, 502, 53–58.

28. Kornberg, A. and Baker, T.A. (2005) DNA replication. 2nd ed. University Science Books, Sausalito, Calif.

29. Frick, D.N. and Richardson, C.C. (2001) DNA primases. Annual review of biochemistry, 70, 39–80.

30. Stratling, W. and Knippers, R. (1973) Function and purification of gene 4 protein of phage T7. Nature, 245, 195–197.

31. Wolfson, J. and Dressler, D. (1972) Regions of single-stranded DNA in the growing points of replicating bacteriophage T7 chromosomes. Proceedings of the National Academy of Sciences of the United States of America, 69, 2682–2686.

32. Richardson, C.C., Romano, L.J., Kolodner, R., LeClerc, J.E., Tamanoi, F., Engler, M.J., Dean, F.B. and Richardson, D.S. (1979) Replication of bacteriophage T7 DNA by purified proteins. Cold Spring Harbor symposia on quantitative biology, 43 Pt 1, 427–440.

33. Lee, S.J., Zhu, B., Hamdan, S.M. and Richardson, C.C. (2010) Mechanism of sequence-specific template binding by the DNA primase of bacteriophage T7. Nucleic acids research, 38, 4372–4383.

34. Corn, J.E., Pease, P.J., Hura, G.L. and Berger, J.M. (2005) Crosstalk between primase subunits can act to regulate primer synthesis in trans. Molecular cell, 20, 391–401.

35. Corn, J.E., Pelton, J.G. and Berger, J.M. (2008) Identification of a DNA primase template tracking site redefines the geometry of primer synthesis. Nature structural & molecular biology, 15, 163–169.

36. Andrilenas, K.K., Penvose, A. and Siggers, T. (2015) Using protein-binding microarrays to study transcription factor specificity: homologs, isoforms and complexes. Briefings in functional genomics, 14, 17–29.

37. Soultanas, P. (2005) The bacterial helicase-primase interaction: a common structural/functional module. Structure, 13, 839–844.

38. Thirlway, J. and Soultanas, P. (2006) In the Bacillus stearothermophilus DnaB-DnaG complex, the activities of the two proteins are modulated by distinct but overlapping networks of residues. Journal of bacteriology, 188, 1534–1539.

39. Naue, N., Beerbaum, M., Bogutzki, A., Schmieder, P. and Curth, U. (2013) The helicase-binding domain of Escherichia coli DnaG primase interacts with the highly conserved C-terminal region of single-stranded DNA-binding protein. Nucleic acids research, 41, 4507–4517.

40. Chintakayala, K., Larson, M.A., Grainger, W.H., Scott, D.J., Griep, M.A., Hinrichs, S.H. and Soultanas, P. (2007) Domain swapping reveals that the C-and N-terminal domains of DnaG and DnaB, respectively, are functional homologues. Molecular microbiology, 63, 1629–1639.

41. Afek, A., Ilic, S., Horton, J., Lukatsky, D.B., Gordan, R. and Akabayov, B. (2018) DNA Sequence Context Controls the Binding and Processivity of the T7 DNA Primase. iScience, 2, 141–147.

42. Ilic, S., Cohen, S., Afek, A., Gordan, R., Lukatsky, D.B. and Akabayov, B. (2019) DNA Sequence Recognition by DNA Primase Using High-Throughput Primase Profiling. J Vis Exp.

43. Balakrishnan, L. and Bambara, R.A. (2013) Okazaki fragment metabolism. Cold Spring Harbor perspectives in biology, 5.

44. Ward, J.H.J. (1963) Hierarchical Grouping to Optimize an Objective Function. J Am Stat Assoc, 58, 236–244.

45. Chen, W., Lei, T.Y., Jin, D.C., Lin, H. and Chou, K.C. (2014) PseKNC: a flexible web server for generating pseudo K-tuple nucleotide composition. Anal Biochem, 456, 53–60.

46. Dao, F.Y., Lv, H., Wang, F. and Ding, H. (2018) Recent Advances on the Machine Learning Methods in Identifying DNA Replication Origins in Eukaryotic Genomics. Front Genet, 9, 613.

47. Tibshirani, R. (1996) Regression Shrinkage and Selection via the lasso. Journal of the Royal Statistical Society, 58, 267–288.

48. Romano, L.J. and Richardson, C.C. (1979) Characterization of the ribonucleic acid primers and the deoxyribonucleic acid product synthesized by the DNA polymerase and gene 4 protein of bacteriophage T7. The Journal of biological chemistry, 254, 10483–10489.

