## Supplementary material for "Reconfiguring primase DNA-recognition sequences by using a data-driven approach": Suplemental Figures and Tables

### METHODS

**Materials**. All chemical reagents were of molecular biology grade and were obtained from Sigma. ATP and CTP were purchased from Roche Molecular Biochemicals.

**Protein overexpression and purification. The** T7 primase domain (residues 1–271) was overexpressed and purified as previously described (1).

**Design of the DNA library.** The analysis was based on previously collected data (2,3), specifically, on 25,220 DNA sequences that include the T7 DNA-primase recognition sequence (5'-GTC-3'). The general pattern of each sequence was 5'-$(N)_{17}$-GTC-$(N)_{16}$-GTCTTGATTCGCTTGACGCTGCTG-3', where $(N)_{17}$ and $(N)_{16}$ represent the variable regions flanking the GTC recognition site. The above data Ωset contained accurate binding scores for T7 primase to each DNA sequence, obtained by PBMs as described previously(3). Data acquisition was performed using a GenePix 4400A scanner (Molecular Devices), and data was analyzed using custom scripts to obtain fluorescence intensities for all sequences represented on the array.

**Data preprocessing.** Each PBM consisted of 5,076 unique sequences and 25,220 samples, 6 repetitions per sequence, and overall 151,320 samples (instances). All scripts were written in Python (Python Software Foundation, version 3.7, http://www.python.org), Scikit learn (4), and the software PyCharm (community edition, https://www.jetbrains.com/pycharm/). The source code for the machine learning algorithms is available in the Github repository (https://github.com/csbarak?tab=repositories). This git repository also contains the data used for the analysis.

By extracting the coefficient of variation (5) for scores associated with each sequence (6 repetitions), we observed that the stronger the score, the more stable the coefficient of variation (Supplementary Figure S1). Finally, each sequence's score was determined as its median score.

It was necessary for the stability evaluation to account for the different binding score ranges; thus, in order to eliminate the different scales of the standard deviation, we evaluated the binding score stability of each sequence by using the coefficient of determination (COD, Eq. 1):

$$COD(x) = \frac{\sigma(x)}{\mu(x)} \qquad (1)$$

where $x$ – is a set of binding score repeats for a specific sequence; σ – is the standard deviation of that sequence; and μ – is the mean value of $x$.

**Method for sequence-based feature extraction.** We have tried linear, quadratic and root weighting of the kmers according to their distance from the GTC. E.g., while the 3-mer ‘ACA’ appears twice in the sequence ‘ACATGTCACAT’, the weighted linear count of ‘ACA’ would multiply its distance plus 1, from the kernel (GTC). However, this approach did not improve the performance of the model. While proper usage of the mer's location might lead to different results, using advanced algorithms to produce a more complex connection between features would limit our work's explainability and further exploration of the mer's effect. The kmer counts were normalized by the length of the sequence to increase the generalization ability of our technique and prevent bias.

**Principal component analysis.** PCA is commonly used to reduce dimensions of datasets by de-correlating the features and extracting the linear combinations that hold the greatest variance. Thus, non-informative features are dropped, and the remaining features consist of highly variant linear combinations (principal components) of the original features. We used PCA on overlapping K-mer count instances so as to visualize the projected distribution of binding scores upon the three most significant principal components. Features were obtained by counting every combination of dimers, trimers, and tetramers in the DNA sequence (K-mer, Supplementary Figure S2). Different K values were used for the K-mer feature extraction, and all experiments resulted in a clear 5-cluster construct for the entire dataset. To compare data between clusters, we applied MinMax normalization (Eq. 2) and colored each instance according to its relative strength.

$$y_i' = \frac{y_i - \min(y)}{\max(y) - \min(y)} \qquad (2)$$

where y = binding scores of the entire dataset, $y_i$ = the i$^{th}$ binding score.

**Conversion of the categorical DNA variables.** The DNA data was converted to an array of integers by OHE, a process in which each nucleotide is represented by the following scheme: (A=[1000], C=[0100], G=[0010], T=[0001]). The N × 4 matrix represents every DNA oligo, and is used as input for both the Kmeans model and the WD-based hierarchical clustering model[17].

**Kmeans.** In the initial step of Kmeans, the distances of the sequence vectors in the training set from randomly located centroids are measured, with the number of centers (K) being considered as a hyper parameter. Then, the distance of every sequence from the centroid is computed using the Euclidean distance ($d(x) = \min_{j=0,1,\ldots,K} \| x - \mu_j \|_{l2}$). For the optimization step, each centroid's position ($\mu_j$) is moved to its own cluster's geometric mean. This process is repeated until a stop condition is met, which is usually determined by an improvement in the loss function. The loss function of the $i^{th}$ iteration is the sum of the distances between all instances and their matching centroids (Eq. 3):

$$L_i(X, \mu_i) = \sum_{x \in X} d(x) \qquad (3)$$

where $i$ is the iteration number, $X$ denotes the entire data matrix, $x$ represents an OHE vector, and $\mu_i$ represents the set of centroids at the beginning of the $i^{th}$ iteration.

An optimized model is obtained when the difference in the value of the loss function between consecutive iterations is small enough (typically $10^{-4}$) or the maximal amount of iterations has been reached.

**Hierarchical clustering.** Ward's criterion is used to determine which clusters should be merged by creating new data partitions in such a way that the sum of cluster variances of the newly offered partitions is kept low; in our case, it amounts to the smallest number of nucleotide changes between same-cluster sequences. Since the sum of the squared errors is minimized when each "word" acts as its own cluster, the common way to choose the number of clusters $K$ is to choose the $K$ that maximizes the WD gap. Using this method, we can extract both $K$ and the evolutionary stages of each cluster. WD calculates the similarity of two clusters $(C_a, C_b)$ as the normalized distance of their corresponding cluster means ($\mu_a, \mu_b$, Eq. 4):

$$WD(C_a, C_b) = \frac{|\mu_a - \mu_b|_{l2}^2}{|C_a||C_b|} \quad (4)$$

The first step of the method initiates a cluster for each instance, and the second seeks the two most similar clusters in terms of WD. When found, these two clusters are united, and the second step is repeated until only one cluster remains.

**Supervised learning – linear regression with L1 regularization (Lasso).** The Lasso algorithm performs liner regression under L1 regularization. Its output is a closed form equation that is generated under the constraint of having the smallest number of variables as possible. The algorithm complies with this constraint by applying a penalty for each variable taken into account in the closed form equation. Simple linear regression uses a weighted combination of features to generate a prediction based on (Eq. 5):

$$Y = \sum_{i=1,2,\dots,p} w_i x_i + b \quad (5)$$

where $x_i$ is the $i^{th}$ feature chosen from $p$ features, while $w_i$ and $b$ are the learned weights (usually found by minimizing the mean square error over the training set) and the learned bias, respectively. While a simple linear model uses the entire set of features, Lasso applies a loss function on the number of features. Moreover, compared to L2 regularization, L1 regularization facilitates the zeroing out of features rather than minimizing their weights, leading to the selection of a smart subset of features. Using Lasso on our data required two preprocessing stages; the first was extracting K-mer counts for obtaining a simple and general solution, and the second was applying a square root on the binding scores to better match their values for linear regression. The MinMax-wise normalized scores yielded a cross-validated result with a mean absolute error (MAE) value of 0.10, calculated using (Eq. 6):

$$\text{MAE } E(X) = \sum p_{x_i} * x_i \quad (6)$$

where $x_i$ is the MAE of bin $i$ of the bins obtained by Kmeans, and $p_{x_i}$ is the percentage of samples in that bin out of the entire data set.

We evaluated the results with MAE, and obtained the expected error in terms of a weighted MAE, where the weights refer to the percentage of clustered sequences (Eq. 7).

$WMAE_{primo} = \sum_{i=0}^{4} \frac{|C_i|}{|dataset|} MAE_{C_i}$ (7)

where $C_i$ is the $i^{th}$ cluster, $|C_i|$ is the number of sequences belonging to the $i^{th}$ cluster, $|dataset|$ is the size of the entire dataset and $MAE_{C_i}$ is the mean absolute error of the $i^{th}$ cluster.

Our main goal was to develop a predictive model with as small an error as possible, while maintaining model explainability and simplicity. Examining the results of different regression models (Table 3), we see that the smallest error was achieved using XGBoost, yet the difference between the errors of XGBoost and those of Lasso is about 0.5% MAE. In contrast to the decision-tree-based XGBoost, Lasso generates a closed predictive equation (i.e., $score = \alpha_0 + \alpha_1 MER_1 + \alpha_2 MER_2$ ...), and combined with Lasso's L1 regularization, it constrains the number of features and the coefficients needed for the prediction. In addition, in contrast to support-vector-machine (SVM)-based models, Lasso enables limiting the coefficients to positive values, which could lead to a meaningful K-mer addition approach. Lastly, with Lasso the bias can be neutralized, meaning that the prediction is dependent solely on the K-mer count. Increasing the bias further enables a decrease in the variance and therefore a precise prediction.

**Table 3. Comparative analysis of different regressors, where K = 3, with clustering application.**

| Model | KNN | RBF-SVM | Linear-SVM | RF | XGBOOST | LASSO |
|---|---|---|---|---|---|---|
| MAE | 0.096 | 0.093 | 0.091 | 0.094 | 0.088 | 0.093 |

In summary, in this study, we chose to use Lasso, since it provides good performance and a closed predictive expression that is short and (intentionally) consists of non-negative coefficients. Other regression models also generated an expected error that was less than 10% MAE (Table 3), meaning that the data collection and preprocessing techniques were highly informative regarding the researched binding score.

**Oligoribonucleotide synthesis assay.** Synthesis of oligoribonucleotides by DNA primase was performed as described previously[14]. The reaction mixture contained 5 µM DNA sequences generated by our machine-learning prediction algorithms described above, 1 mM ATP, 1 mM [α-$^{32}$P]ATP, and T7 primase in a buffer containing 40 mM Tris-HCl, pH 7.5, 10 mM $MgCl_2$, 10 mM DTT, and 50 mM potassium glutamate. After incubation at room temperature for 10 min, the reaction was terminated by adding an equal volume of sequencing buffer containing 98% formamide, 0.1% bromophenol blue, and 20 mM EDTA. The samples were loaded onto 25% polyacrylamide sequencing gel containing 3 M urea and visualized using autoradiography.

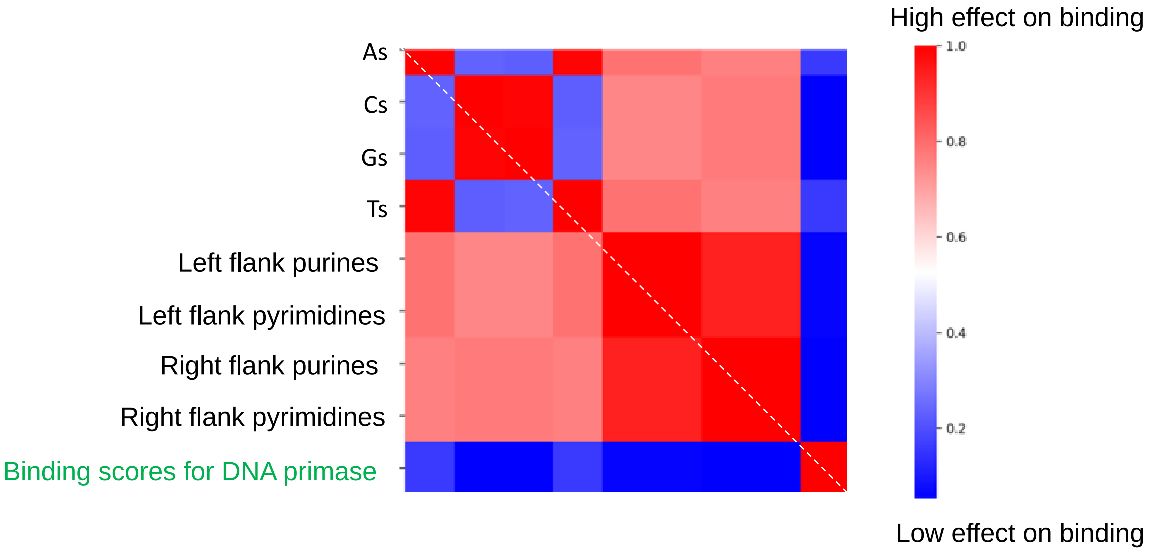


**Figure S1. Diagonal correlation matrix of features representing oligonucleotide composition.** This correlation presents the insignificant effect of hand-crafted features on the binding score of T7 primase. These features were calculated from the sequence of oligonucleotides in the DNA-protein microarray (PBM). Binding score of T7 primase was determined by PBM.

*K-mers for GTCCC*

| *k* | *K-mers* |
|---|---|
| 1 | G, T, C |
| 2 | GT, TC, CC |
| 3 | GTC, TCC, CCC |

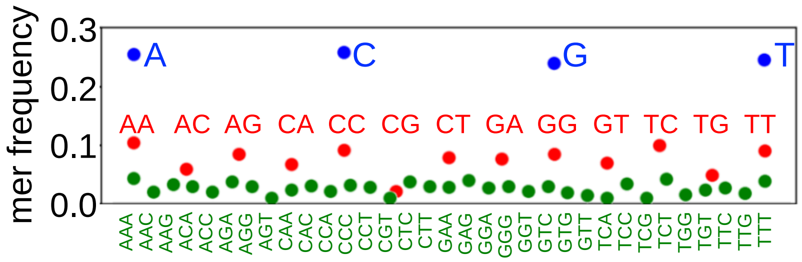


**Figure S2. Illustration of *k-mer* features.** Left: The term *k-mer* refers to all of a sequence's subsequences of length *k*, such that the sequence AGAT would have four monomer (A, G, A, and T), three 2-mers (AG, GA, AT), two 3-mers (AGA and GAT). Right: Higher *k* number is characterized with low frequency of occurrences on the genome.

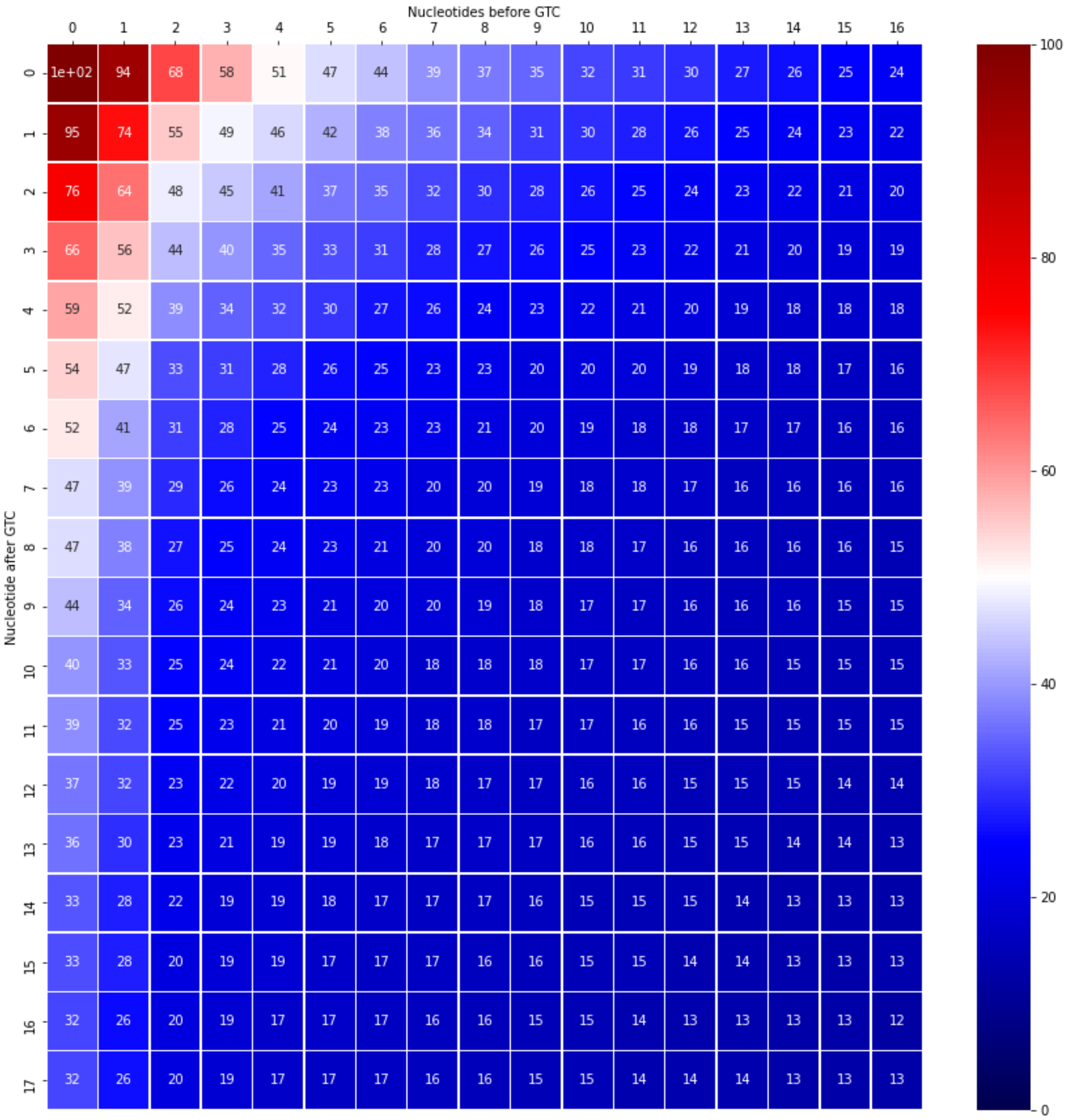


**Figure S3.** Effect of nucleotide position outside the ‘GTC’ on primase binding. Gradient boosting machine (GBM) was trained on a different sliding "window" of nucleotides' positions before and after the labeled GTC-containing sequences. The model is represented by the MAE score on the test data.

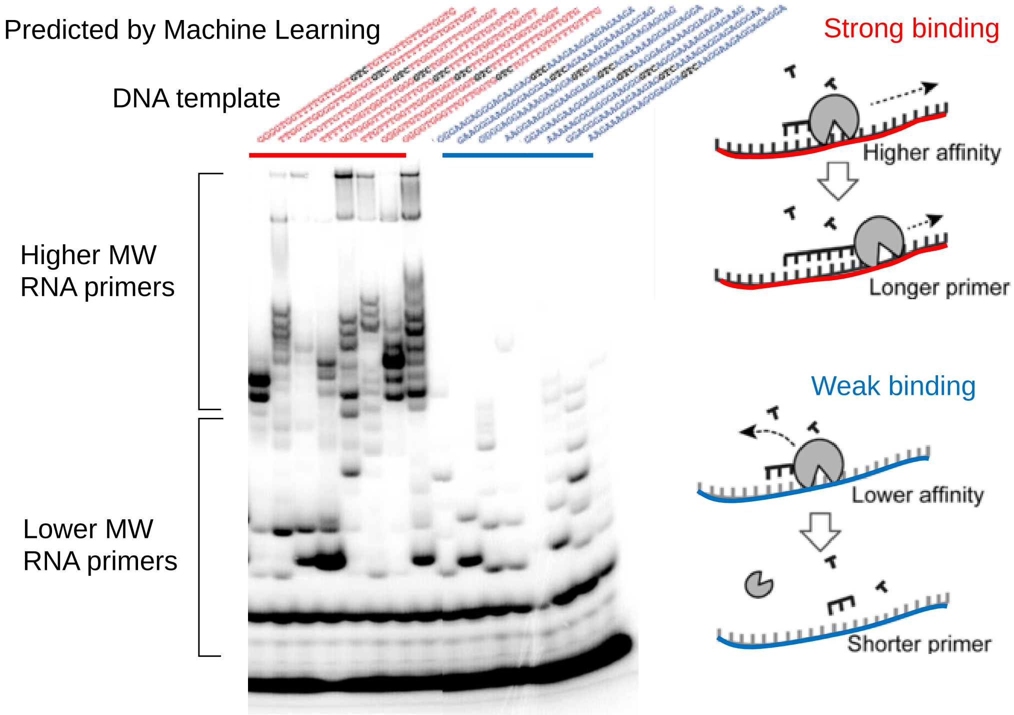


**Figure S4. Template-directed RNA primer synthesis catalyzed by the T7 DNA primase**. Oligonucleotide synthesis by the T7 DNA primase. The reactions contained oligonucleotides with the primase recognition sequence as indicated, and 32P-γ-ATP, ATP, CTP, UTP, and GTP in the standard reaction mixture. After incubation, the radioactive products were analyzed by electrophoresis through a 25% polyacrylamide gel containing 7 M urea, and visualized using autoradiography. Longer RNA primers were formed on DNA templates that were predicted to have higher binding affinity to the T7 primase.

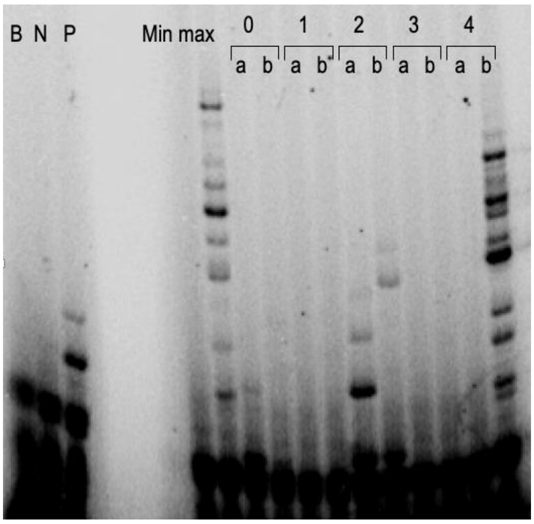


**Figure S5. RNA primer synthesis by T7 DNA primase on selected DNA templates from different clusters obtained by unsupervised learning.** Oligonucleotide synthesis by the T7 DNA primase was performed as described in Supplementary Figure S4. The reactions contained selected oligonucleotides sequences from different clusters obtained by unsupervised analysis (Figure 3).

**Table S1. Test data**

| Sequence | Empirical results | Predicted results |
| --- | --- | --- |
| AAAAAGGGAGGGAAGGGGTCAGGGAAAAGAGAGAAG | 2016 | 0.358457 |
| AAGAAAGGAAGGGAGGAGTCAAGGAAGAGGAGAGGA | 1956.5 | 0.232119 |
| AAGGAAGGGGAAGGAGAGTCAGAAAAAGGAGGAGGA | 2570 | 0.294887 |
| GAAGGGAAGGGGAGGAAGTCAGAAAAGAAAGAGGAG | 2365 | 0.292978 |
| GGAGAAGAAGGAGGAGAGTCAAGGAGAAAAGGAGGA | 1650 | 0.244383 |
| GGAGGGAAAGAGAAGAGGTCAAAAGAGGAGAGGGAA | 1904.5 | 0.316637 |
| GGGAAGAGGGAGAAGAGGTCAAAGAAGGAGAGAAGA | 1859 | 0.292191 |
| GGGGAGGAAAAGAAGGAGTCAGAAGAAGAAGAGGAG | 1895 | 0.282706 |
| GGGGTGGGTTGTTGGTGGTCTGTTTGTGTTTGTTTG | 48919.5 | 0.784515 |
| GGGGTGGTTTTGTTGGTGTCTGTTGTTGTTGTGGTG | 49672 | 0.804615 |
| GGGGTGTGGTGGGTGGTGTCTTTTTTTTTGGTTGTG | 48331.5 | 0.815105 |
| GGTGGGTTTGTGTTGTGGTCTTTTGTGGTGTGGGTT | 45403 | 0.796791 |
| GGTGTTGTTGGTGGTGTGTCTTGGTGTTTTGGTGGT | 41314.5 | 0.822484 |
| TTGGTTGGGGTTGGTGTGTCTGTTTTTGGTGGTGGT | 43744 | 0.745619 |
| TTGTTTGGTTGGGTGGTGTCTTGGTTGTGGTGTGGT | 41522 | 0.79126 |
| TTTTTGGGTGGGTTGGGGTCTGGGTTTTGTGTGTTG | 51845.5 | 0.658822 |

**Table S2. DNA sequences from generative algorithm**

| Cluster | Sequence | Predicted results |
| --- | --- | --- |
| max | CCACCCCAAAAAACCCCGTCAAAACCCCAAAAACCA | 1 |
| min | GACGAAGACGACGAAGAGTCCGAGGAAGCAGACGAA | 0 |
| 0 | CCCAAAAAACCCCAAAAGTCTCCACCAACCCCAAAA | 0.471915441 |
| 0 | CCAAAACAAACCCAACAGTCACCACCCCACCCTAAA | 0.629928763 |
| 1 | AAGGAAGGAGAAGAGAAGTCGAGGGAGAGCGAGGAA | 0.136521961 |
| 1 | GGAGAAGAGGAGGAGGTGTCAGAAGAAAAAGAAAGG | 0.241974161 |
| 2 | CCTCCCTTTTTTTTTTTGTCCTCTCCTCCTTTCCCC | 0.316487367 |
| 2 | TTCCACCACTCCATTCTGTCAACGTATTCTTCACCC | 0.50895981 |
| 3 | CTTCGAAGCAACCAAAGGTCGCAAGTTGAATAAGAC | 0.468099743 |
| 3 | CGATGCTGTTCCGTTTGGTCAACTAAAGACCATGAT | 0.502488194 |
| 4 | TTTTTTTTTTTGGGGGGGTCGGGTTTGGGGTGGGGT | 0.669837793 |
| 4 | TTGTGTGGGGTCTTGTGGTCTTTGTGTGTTGGGTGT | 0.805995072 |

**REFERENCES**


1. Zhu, B., Lee, S.J. and Richardson, C.C. (2009) An in trans interaction at the interface of the helicase and primase domains of the hexameric gene 4 protein of bacteriophage T7 modulates their activities. *The Journal of biological chemistry*, **284**, 23842-23851.
2. Ilic, S., Cohen, S., Afek, A., Gordan, R., Lukatsky, D.B. and Akabayov, B. (2019) DNA Sequence Recognition by DNA Primase Using High-Throughput Primase Profiling. *J Vis Exp*.
3. Afek, A., Ilic, S., Horton, J., Lukatsky, D.B., Gordan, R. and Akabayov, B. (2018) DNA Sequence Context Controls the Binding and Processivity of the T7 DNA Primase. *iScience*, **2**, 141-147.
4. Pedregosa, F., Varoquaux, G., Gramfort, A., Michel, V., Thirion, B., Grisel, O., Blondel, M., Prettenhofer, P., Weiss, O., Dubourg, V. *et al.* (2011) Scikit-learn: Machine Learning in Python. *Journal of Machine Learning Research*, **12**, 2825-2830.
5. Brown, C.E. (1998) *Coefficient of Variation*. Springer, Berlin, Heidelberg.
